# *Mus musculus* populations in Western Australia lack VKORC1 mutations conferring resistance to first generation anticoagulant rodenticides: Implications for conservation and biosecurity

**DOI:** 10.1101/2020.07.06.189282

**Authors:** Bridget J.M.L. Duncan, Annette Koenders, Quinton Burnham, Michael T. Lohr

**Affiliations:** Centre for Ecosystem Management, School of Science, Edith Cowan University, 270 Joondalup Drive, Joondalup WA 6027, Australia

## Abstract

**Background:** Humans routinely attempt to manage pest rodent populations with anticoagulant rodenticides (ARs). We require information on resistance to ARs within rodent populations to have effective eradication programs that minimise exposure in non-target species. Mutations to the VKORC1 gene have been shown to confer resistance in rodents with high proportions of resistance in mice found in all European populations tested. We screened mutations in *Mus musculus* within Western Australia, by sampling populations from the capital city (Perth) and a remote island (Browse Island). These are the first Australian mouse populations screened for resistance using this method. Additionally, the mitochondrial D-loop of house mice was sequenced to explore population genetic structure, identify the origin of Western Australian mice, and to elucidate whether resistance was linked to certain haplotypes.

**Results:** No resistance-related VKORC1 mutations were detected in either house mouse population. A genetic introgression in the intronic sequence of the VKORC1 gene of Browse Island house mouse was detected which is thought to have originated through hybridisation with the Algerian mouse (*Mus spretus*). Analysis of the mitochondrial D-loop reported two haplotypes in the house mouse population of Perth, and two haplotypes in the population of Browse Island.

**Conclusions:** Both house mouse populations exhibited no genetic resistance to ARs, in spite of free use of ARs in Western Australia. Therefore weaker anticoagulant rodenticides can be employed in pest control and eradication attempts, which will result in reduced negative impacts on non-target species. Biosecurity measures must be in place to avoid introduction of resistant house mice, and new house mouse subspecies to Western Australia.

## Introduction

Commensal rodents are resident on all land masses (except Antarctica)[1,2] and are expanding their geographical distributions, which is indicative of a high adaptability to challenging environmental conditions and ability to exploit resources provided by humans.[3] These rodents are considered pests, and poisons are the most commonly used method for control and eradication programs.[4] Due to their efficacy, anticoagulant rodenticides (ARs) have been used to carry out eradications on over 300 islands with a 90% success rate.[5,6] Anticoagulant rodenticides can be subdivided into first generation (FGARs) or second generation (SGARs), according to when they were first synthesised and commercialised.[7] In 1958, the first evidence of resistance to ARs was discovered in *Mus musculus* populations, which were resistant to warfarin and diphacinone.[8] Second generation compounds with greater acute toxicities than FGARs were developed in response, resulting in mortality after a single feed at a lower concentration.[7,9,10]

Resistance-linked mutations within rodent species can arise independently and be positively selected in populations that are exposed to ARs, or can be acquired via introgression with other species, for example, by hybridisation between *M. musculus* and the Algerian mouse (*Mus spretus*).[11–13] Resistance to ARs in *M. musculus* populations has been linked to single nucleotide polymorphisms in subunit 1 of the *Vitamin K epOxide Reductase Complex* (VKORC1) which is involved in the synthesis of vitamin K and is important in blot clotting[14] and bone formation.[15] However, mutations in the VKORC1 gene should be selected against in the absence of exposure to rodenticides due to negative pleiotropic effects.[14–17] Extensive testing of resistance to SGARs in *M. musculus* in Europe, Canada and some in Australia via feeding trials, blood clotting response tests, and/or genetic screening has uncovered high prevalence of SGAR resistance.[11,18,19] For example, every European population tested (100 from 6 countries) had levels of VKORC1-related resistance ranging from 70 to 100%.[11,13,19–21]

It is vital to understand levels of resistance in rodent populations so that appropriate chemicals can be applied at a suitable rate for management purposes, as chemical residues of ARs have been detected in a wide variety of non-target species, in both terrestrial and aquatic environments. The presence of these chemicals outside of their target species can result directly from the rodenticide, or indirectly via consumption of poisoned rodents as persistence of rodenticides[22–24] results in liver retention and ultimately biomagnification and bioaccumulation in food webs.[9]

There is scarce information regarding the presence and distribution of resistance in rodent populations of Australia, and a complete absence of information for *M. musculus* populations on the Australian mainland.[25,26] In Western Australia, neither rats nor mice have been tested to quantify the levels of resistance to ARs despite the pervasive use of poisons by government agencies and the public, whose use of both FGARs and SGARs is unmonitored and unregulated.[9]

As ARs are the most commonly used class of chemicals to control both mainland and island rodent populations in Australia, it is critical to evaluate the levels of resistance in rodent populations in order to reduce the impact on non-target species.[9] For this reason, the aim of this study was to investigate whether VKORC1 mutations that have been linked to AR resistance are present within populations of *M. musculus* in the urban area of Perth (the largest city in Western Australia). As a comparison, the same investigation was undertaken for a population of south-east Asian house mouse (*M. musculus castaneus*) from Browse Island off the coast of Western Australia where ARs have not been used. For both populations, analyses of the mtDNA D-loop region were conducted to investigate whether resistance (if present) is correlated with genetic lineages within these species.

## Materials and methods

### Sample collection and preparation

Western house mice (*Mus musculus domesticus*) from the Perth metropolitan area (Western Australia) were collected by use of traps or other methods not involving ARs (n=34); confirmation that death was not due to ARs was derived from the absence of internal haemorrhaging noted during dissection. Mice were stored at −20°C at Edith Cowan University until liver extraction, and livers were then stored in 100% ethanol at −8°C until DNA extraction. Liver samples of south-east Asian house mouse (*Mus musculus castaneus*) that had been collected using snap traps from Browse Island were donated by the Department of Biodiversity, Conservation and Attractions of Western Australia (n=15). Liver samples from these mice were preserved in salt saturated dimethyl sulfoxide and stored at −8°C at Edith Cowan University until DNA extraction.

### Sensitivity analysis

We carried out a sensitivity analysis to ascertain the likelihood of detecting mutations in our sample of 34 mice under different proportions of mutation in the Perth population. Using a binomial distribution, we tested proportions of resistant individuals in the population ranging from 70%[11] (the lowest percentage of resistance detected in the literature) down to 10%. The probability of not collecting any mice with resistance in our sample size of 34 was well below 0.000001 at 70% and 0.028 at 10% in the population (Table 1). In fact, even at the lowest resistance rate tested, we could have expected to collect at least 3 resistant mice, and for a population with 70% resistance that number rises to 24 mice.

**Table 1.** Binomial sensitivity analysis.

|  | Proportion of resistant mice in population (%) |  |  |  |
| --- | --- | --- | --- | --- |
|  | 10 | 30 | 70 | 95 |
| Probability of finding no resistant mice | 0.028 | <0.0001 | <0.000001 | <0.000001 |
| Mode (resistant mice detected) | 3 | 10 | 24 | 33 |

Finding a resistant mouse was taken as a success. The analysis was based on the Perth population with a sample size of 34.

### DNA extraction and PCR

Prior to DNA extraction, the liver was washed in Milli-Q water for 5 minutes on a rocking platform at room temperature, then the water was drained and replaced, and the process repeated under the same conditions for 10 minutes. Genomic DNA was extracted from the liver using a DNeasy Blood & Tissue kit following the manufacturer’s instructions (Qiagen). Concentration and purity of the DNA were measured using a NanoDrop (Thermo Fisher Scientific), and then concentration of DNA in all samples was standardised to 10 ng/μL by dilution with Milli-Q water unless it was below 10 ng/μL.

Polymerase chain reaction was used to amplify the three exons of the VKORC1 gene region^21^ and the complete mitochondrial D-loop and flanking regions^32^ using primer sets from the sources indicated. Reactions included 20 ng of DNA template, 12.5 μL of Platinum II HotStart (VKORC1) or Platinum HotStart (D-loop) master mix (Thermo Fisher Scientific), 0.5 μL of both forward and reverse primers made up to 25 μL with DNA-free H_2_O. Cycling conditions for PCR are shown in Table 2. Successfully amplified PCR products were purified and sequenced at the Australian Genome Research Facility in Perth, Western Australia.

**Table 2.** Adapted PCR parameters for VKORC1 and D-loop amplification.[11]

| Gene region | Master Mix | Initial denaturation | Denaturation | Annealing | Extension | Final extension |
| --- | --- | --- | --- | --- | --- | --- |
| VKORC1 | Platinum HotStart | 2 min @ 94°C | 30 sec @ 94°C | 30 sec @ 58°C | 110 sec @ 72°C | 10 min @ 72°C |
|  | Platinum II<br>HotStart | 2 min @ 94°C | 30 sec @<br>94°C | 30 sec @<br>60°C | 30 sec @<br>72°C | 10 min @<br>72°C |
| D-loop | Platinum<br>HotStart | 2 min @ 94°C | 30 sec @<br>94°C | 30 sec @<br>55°C | 1 min @<br>72°C | 10 min @<br>72°C |

All conditions included 40 cycles.

### Sequence construction

Sequence chromatograms were manually checked and edited using FinchTV (Geospiza) and aligned in Geneious Prime 2019.0.4 with multiple reference *Mus musculus* and *Mus spretus* sequences downloaded from GenBank to confirm (sub)species identification, identify haplotypes, and detect VKORC1 SNPs. Reference sequences for the VKORC1 gene included wild type (susceptible to ARs) *M. m. domesticus*, AR resistant *M. m. domesticus*, the Algerian mouse *M. spretus*, and the hybridised (and resistant to ARs) *M. m. domesticus/ M.spretus*.

### Haplotype network construction

The D-loop sequences of Perth metropolitan mice and Browse Island specimens were aligned in Geneious with 578 *M. m. domesticus* D-loop sequences and 246 *M. m. castaneus* D-loop sequences, respectively. Non-unique sequences from the same country were filtered out from both datasets, and sequence length was standardised. Haplotype networks were created from the aligned datasets using the minimum spanning network method in PopART (Population Analysis with Reticulate Trees).[27]

### Ethical approval

All protocols were approved by the Edith Cowan University Animal Ethics Committee (permit number 21776).

## Results

### VKORC1 mutations, insertions and deletions

VKORC1 amplification covered three exons. Exons 1 and 2 were amplified using one primer set and resulted in complete sequences for 31 mice and incomplete sequences in 21 mice. Exon 3 resulted in 26 complete sequences and one incomplete. In total, 15 mice from Browse Island and 34 mice from Perth were sequenced, in approximately equal sex ratio.

Sequences from *M. m. castaneus* presented two noteworthy features: all mice carried two silent single nucleotide polymorphisms in exon 1 at amino acid positions 10 and 37, and all mice had a 68 base pair insertion at nucleotide position 1454, located in the intron following exon 2. When compared to published sequences it was found that the insertion (12 mice) was 100% homologous to a section of DNA in *M. spretus* as well as certain *M. m. domesticus* that hybridised with the Algerian mouse. The insertion was not identical in all individuals sequenced from the island: the remaining three individuals differed from the published sequence by a single identical point mutation. None of the mutations linked to the *M. spretus* resistant allele were detected in any of the Browse Island mice, despite all displaying the intragenic sequence. Furthermore, all 15 Browse Island samples reported an eight bp deletion at position 1143 that is shared with the published sequences previously mentioned. One mouse from Browse Island displayed a seven bp and an 11 bp insertion at nucleotide position 939 and 1266 respectively, which were not exhibited in any of the published VKORC1 sequences. The VKORC1 insertion discovered on Browse Island is here defined as an introgression in agreement with previous studies that have used simulation and statistical methods to reach this conclusion.[28–30]

In contrast, no mutations (associated with AR resistance or not) were detected in the coding regions of the VKORC1 gene in western house mice (*Mus musculus domesticus*) from the Perth metropolitan area.

### *Mus musculus* D-loop haplotypes in Western Australia

Mitochondrial D-loop sequences were obtained for all house mice in this study (n=49), with sequenced PCR products between 900 and 1100 base pairs. The Perth mouse population had two haplotypes, with the most common being 100% homologous to published sequences of *M. m. domesticus* from Australia. One mouse from the metropolitan area displayed a different haplotype, which did not match any published D-loop sequences; this mouse was most closely related to haplotypes from Australia and The Netherlands, but differed from them by a single base pair change at position 718. The haplotype network (Fig 1) groups Perth mice with haplotypes from West Europe, New Zealand, and the British Isles, as well as Southeast Asia, Africa and Oceanic Islands. The western house mouse from Perth with a unique haplotype was most closely related to *M. m. domesticus* from the Middle East, Cyprus, and Europe.

**Figure 1.**
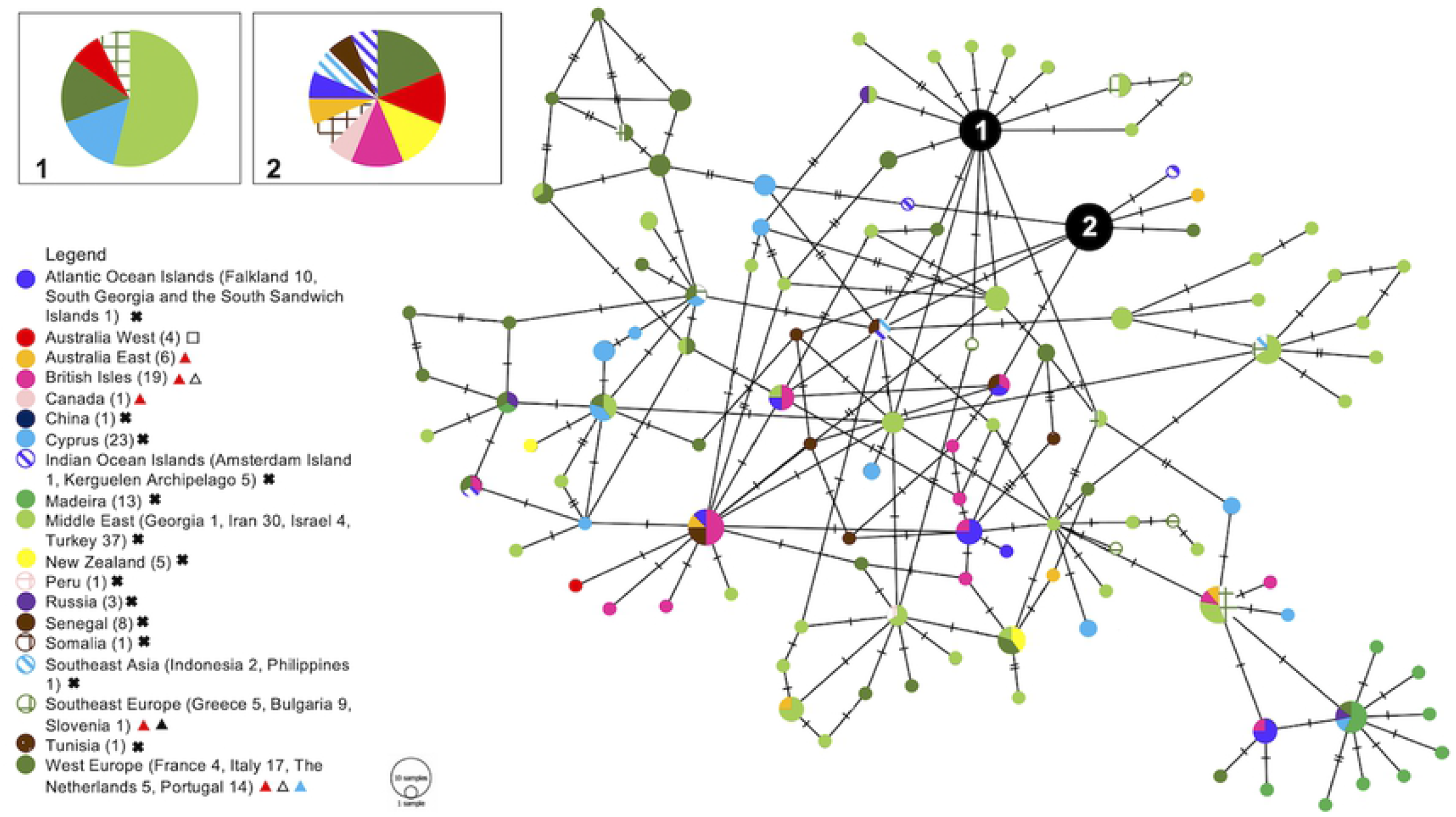
Minimum spanning haplotype network from *Mus musculus domesticus* D-loop data. The numbers in the legend are representative of the number of unique haplotypes included in the analysis. The size of each circle indicates the frequencies of the haplotype, and each colour represents the geographical origin of the sequences. The numbered nodes in the haplotype network correspond to the numbered inset boxes. A black cross in legend indicates that M. m. domesticus in that area has not been tested for anticoagulant rodenticide (AR) resistance.[31] A white square indicates that M. m. domesticus has been tested for Vkorc1 mutations related to AR resistance and was found to be negative (this study). A red triangle indicates that M. m. domesticus has been tested for resistance through a lethal feeding period test and was found to be resistant to ARs.[5,8,20,25,26] A white triangle indicates that M. m. domesticus has displayed Vkorc1 mutations related to AR resistance.[20] A black triangle indicates that a blood clotting response test has been performed and M. m. domesticus was found to be resistant to ARs.[26] A blue triangle indicates that an assay of the VKOR enzyme has been carried out and resistance to ARs was determined.[5]

The D-loop sequences of 15 mice from Browse Island were identical to published *M. m. castaneus* D-loop sequences from Asia and Kenya. One mouse differed from the other haplotype by one base pair at position 101 and was 100% homologous to published *M. m. castaneus* sequences from Asia, Russia and Kenya. On the haplotype network (Fig 2), mitochondrial D-loop sequences of *M. m. castaneus* from Browse Island were grouped with sequences from China, Bangladesh, Japan, Indonesia and Kenya, as well as other Asian countries.

**Fig 2.**
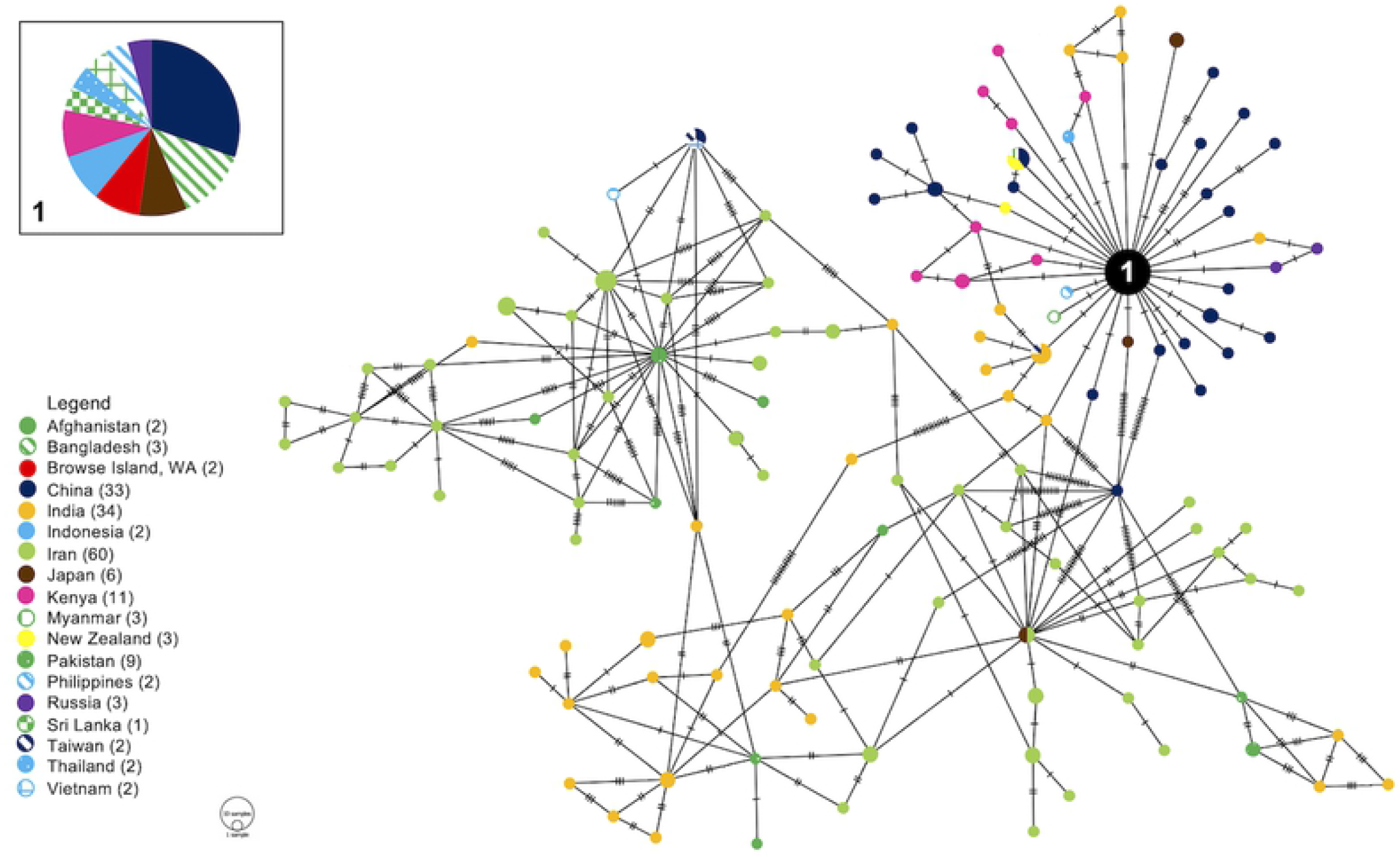
Minimum spanning haplotype network from mitochondrial D-loop data of *Mus musculus castaneus*. The numbers in the legend are representative of the number of unique haplotypes included in the analysis. The size of each circle indicates the frequencies of the haplotype, and each colour represents the geographical origin of the sequences. The numbered node in the haplotype network corresponds to the numbered inset box. Haplotype diversity resulting from deletions and insertions is not represented in PopART (1.7) networks, therefore all Browse Island samples appear in the same haplotype node.

## Discussion

### *Mus musculus domesticus* of Perth lack VKORC1 mutations

Resistance to ARs is usually present in populations of house mice and rats that have been exposed to toxicants, especially when exposure is repeated and involves intensive use of FGARs such as warfarin.[32] ARs have been intensively used by the public and professionals for conservation outcomes, or in agricultural, residential and commercial settings in Western Australia,[9] and professional pest managers have anecdotally observed resistance to the ARs warfarin, coumatetralyl (FGARs) and bromadiolone (SGAR) in Perth house mice (Piggott D, 2019, pers. comm.). Where mouse populations carry resistance, larger quantities or stronger ARs tend to be used, increasing the likelihood of non-target animals being affected directly through feeding on baits and indirectly through feeding on resistant mice that have ingested higher doses of poison (in addition to consuming dead susceptible mice).

Prior to this study, the resistance to warfarin in Australian *Mus musculus* to ARs had been recorded only using no-choice feeding tests on animals from Lord Howe Island (700 km off the coast of New South Wales), and Sydney but not in Western Australian populations.[25,33] In Australia, the *Vkroc1* gene has not previously been sequenced in mice, and therefore the results of this study are the only available information currently on genetically-based resistance to ARs of mice in Australia.

No mutations correlated with resistance to ARs were detected in the VKORC1 gene of *M. m. domesticus* from the Perth metropolitan area in this study. Previous VKORC1 screening in Europe has recorded high frequencies of mutations related to AR resistance ranging 70-100% (Table 3).[11,19,21,28,34] All published research at the time of preparing this manuscript reported resistance-related mutations, therefore the absence of VKORC1 sequence variants in the Perth region is a unique result and poses challenges and opportunities for pest management and biosecurity.

**Table 3.** Anticoagulant resistance conferred by mutations in the VKORC1 gene in *Mus musculus* populations worldwide.

| Location | Number of mice | Resistance (%) | Reference |
| --- | --- | --- | --- |
| France (65 locations) | 266 | 70 | [11] |
| United Kingdom (5 counties) | 53 | 89 | [20] |
| Germany (25 locations) | 352 | 90 | [19] |
| Ireland | 50 | 84 | [21] |
| Switzerland (3 locations) | 48 | 100 | [19] |
| Spain | 29 | 93 | [30] |

We consider six scenarios as likely explanations for the lack of genetic resistance in Perth mice (Table 4). The simplest scenario is a founder effect: the mutations in the VKORC1 gene were never present in the Western Australian mice population. As *M. m. domesticus* was probably introduced in Australia by European settlers in the late 18^th^ Century, the absence of mutations in the exons of VKORC1 might because they were absent from these founder animals.[35,36] The mitochondrial D-loop data supports this, as the low haplotype diversity indicates rapid expansion from a small founding population.[37] ARs were invented in the 1940s, and the first evidence of resistance in rodents dates to 1958.[18] Therefore, the highly conserved coding regions of VKORC1 have been maintained free of mutations because the initial rodent population that colonised Western Australia did not exhibit any.

**Table 4.** Possible scenarios that could have led to the absence of SNPs in the VKORC1 gene of *Mus musculus domesticus* of the Perth metropolitan area, Western Australia.

| Scenario | Description | Likelihood |
| --- | --- | --- |
| Founder effect | Mutations have never been in Western Australia because the founder population did not carry VKORC1 mutations. | Possible |
| SGARs | The use of strong SGARs removed any warfarin resistance (the most common kind of resistance inferred from VKORC1 mutations) from the Western Australian house mouse population. | Very likely |
| Pleiotropy | Defective VKOR enzyme activity posed a negative selection pressure on VKORC1 mutations related to rodenticide resistance. | Not likely |
| Other factors | The arid Australian environment selects for natural tolerance to vitamin K depleted conditions; other unidentified genes (such as cytochrome P450) confer resistance to ARs in Western Australian mice. | Possible |
| Detection failure | As only a subset of the Western Australian house mouse population was tested, the sample could have missed individuals from the population carrying VKORC1 mutations. | Not likely |

Alternatively, western house mice of Western Australia may have lost mutations conferring AR resistance. In this scenario, it may be that heavy use of SGARs has removed them from the population. In Australia, eight ARs are commercially available for rodent control, which are intensively used, and readily available to the public without restrictions.[9] While there is no information available regarding the volume of use of rodenticides in Australia, evidence of secondary poisoning and AR residues in non-target species indicates heavy use of rodenticides in Western Australia, both by professionals and the public.[9] As the most common resistance is exhibited towards the anticoagulant warfarin, the use of stronger SGARs may have removed any individuals carrying mutations related to warfarin resistance from Western Australian populations.

Another explanation could be that a large pleiotropic has removed the alleles. Dysfunctional VKORC1 activity has a cost, as blood clotting requires the enzyme to perform well.[14] Furthermore, vitamin K is essential during foetal bone formation and therefore females with normal VKOR activity are better breeders.[15] VKORC1 mutations are therefore selected against in conditions where rodenticides are not used and rodent populations are not exposed to the poisons;[14–17] however, continuous exposure of house mice to ARs in the Perth Metropolitan area poses a selection pressure to maintain resistant mutations, and therefore it is unlikely that mutations in the VKORC1 gene are absent due to negative pleiotropic effects, and resistant rodents might be consuming exogenous vitamin K to overcome the defective enzyme activity. [17]

It is also possible that Perth mice have AR resistance through other mechanisms (i.e. through mutations in other genes conferring resistance), which would result in there being no selection in favour of VKORC1 mutations (and they may be actively selected against as per the pleiotropy scenario). The absence of resistance-related alleles in the VKORC1 gene does not rule out the presence of resistance in Western Australia as this gene is not the only known mechanism of resistance to anticoagulant rodenticides. Cytochrome P450 2C9 (CYP2C9) metabolises anticoagulant rodenticides, and genetic variants are associated with the need for higher dosages.[26,38] As Cytochrome P450 also detoxifies other pollutants, such as pesticides, the presence of environmental pollutants in general could lead to an increase of cytochrome P450 activity and render rodents pre-adapted to rodenticides.[39] Other unidentified genes could be involved in rodenticide resistance as well, and resistance to certain ARs, such as difenacoum, has been shown to be polygenic.[40,41]

Rodents adapted to living in arid environments are tolerant to anticoagulant rodenticides, an adaptation to a scarcity of available vitamin K. This could be an alternative cause of AR resistance in western house mice of Western Australia, where the climate ranges from Mediterranean in the south (where Perth is located) and transitions through semi-arid to desert further inland.[38,42,43] Until resistance to anticoagulant rodenticides is tested in *M. m. domesticus* of Western Australia through laboratory tests such as feeding trials and blood clot response tests, it cannot be said whether the mice of this region are resistant to ARs even though this study has shown that they do not carry resistance-linked alleles in the VKORC1 gene. Pest managers have anecdotally reported resistance in the Perth region, and therefore it is possible that house mice of Western Australia are resistant.

A final possibility is that mutations in the VKORC1 gene of Perth house mice are present but were not detected. Sample size (34 mice) was adequate to detect resistant mice at levels of resistance in the population to as low as 10% (Table 1). Measures were taken to avoid sample collection bias according to sex (equal number of male and female samples were used), location (a geographical range spanning through the entire Perth metropolitan area was preferred), and sample collection was designed not to underestimate the frequency of VKORC1 mutations (i.e. no mice killed with rodenticides were collected). Furthermore, western house mouse populations in Europe reported high percentages of resistance (between 70% to 100%) with much smaller sample sizes than our study (as few as four mice per location)[11] while all other tests performed on house mouse populations worldwide detected alleles associated with resistance.[11,19–21,30] We therefore hold the possibility of a false negative not to be a concern.

Combinations of the scenarios discussed could be active (potentially in different places and at different times); for example, if the initial colonising population did not have mutations in the VKORC1 gene (scenario founder effect), any mutations arising independently could have been quickly removed from the population by the intensive use of strong anticoagulant rodenticides that killed those individuals (scenario SGARs).

### VKORC1 introgression between *Mus* species

*M. m. castaneus* inhabiting Browse Island had an intragenic insertion in the VKORC1 gene, which was 100% homologous with the GenBank *Mus spretus* reference sequence, as well as some *M. m. domesticus* that have hybridised with *M. spretus*.[30] There is evidence of an ongoing process of hybridisation between the *M. musculus* subspecies, which, combined with the retention of ancestral polymorphism and introgression in the wild, renders it difficult to distinguish between subspecies.[44]

The most researched *M. musculus* subspecies, *M. m. musculus* and *M. m. domesticus*[45] present evidence for introgression in at least 10% of their nuclear genomes, as well as multiple mitochondrial restriction sites.[46,47] When compared to *M. m. musculus* and *M. m. domesticus, M. m. castaneus* is more polymorphic, and its retention of ancestral polymorphisms might be the reason behind the complexity of defining its lineage and identifying introgression,[45] making it difficult to interpret the history of the Browse Island mice.

The intragenic insertion detected in this study in the VKORC1 gene has been previously found to have transferred from *M. spretus* to *M. m. domesticus* in areas of sympatry in Spain and allopatry in Germany, where hybridisation and introgression between allopatric species might have been caused by human interaction that increased the geographic distribution of the *Mus* species.[30] Although there is a selective disadvantage from hybridisation between *M. musculus* and *M. spretus*, including sterile male offspring and the possibility of sterile female offspring depending on the direction of the backcross,[48] the introgression of the *M. spretus* haplotype between distant populations has been possible because it involved an adaptive gene such as VKORC1 and its spread has been driven by the selection pressure posed by the common use of ARs.[29]

Previous studies identified three distinct introgression events between *M. m. domesticus* and *M. spretus*, including an ancient event that occurred more than 2000 years ago prior to the colonisation of Europe by *M. musculus.[28]* The other more recent introgression events are dated around 50 to 60 years ago, since the introduction of ARs.[30] Introgression in the VKORC1 gene was present but in low frequencies and it was therefore considered far from becoming fixed and the details were not reported.[29] The 68 bp insertion detected in the present study was present in all Browse Island mice so it can be concluded that the introgression event was likely to be fixed in this small population due to founder effect and genetic bottlenecks. Therefore the region of the VKORC1 gene from *M. spretus* that has been detected in the *M. m. castaneus* sample from Browse Island is considered a comet allele[47] that has migrated from one evolutionary lineage to another and its frequency has increased to reach fixation.

The D-loop places the mice on Browse Island most closely to mice from China, Indonesia, and other Asian countries, but as no other studies have sequenced the VKORC1 gene region in this subspecies or attempted to identify introgression in different *Mus* species from Asian countries, we cannot make inferences on the process and location of the hybridisation between the two *Mus* species. Less is known about hybridisation involving *M. m. castaneus* than the European subspecies;[49] additionally, introgression in the VKORC1 gene between *M. spretus* and *M. m. castaneus* has not been described previously, possibly because the two taxa are allopatric.

The length of the insertion in the VKORC1 gene (68bp), could indicate an ancient introgression event (>2000 years ago) characterised by continued backcrossing, drift and recombination;[28] however the insertion is not fragmented, and is identical across most individuals from Browse Island and the reference sequences from GenBank. The fact that the region is highly conserved is evidence of its adaptive value as selection would maintain it over time, but this affects the length-based method used to determine the age of the hybridisation. The location of the insertion on the gene could influence the way it is conserved. Noncoding intron regions are usually conserved when the exons are alternatively spliced.[50] In humans, the VKORC1 gene is alternatively spliced, and three isoforms have been described, while the VKORC1 of *M. musculus* has a single described isoform and two potential isoforms that have been computationally mapped.[51] If the VKORC1 gene of *Mus musculus* is indeed alternatively spliced, the possibility of the intron being conserved is much higher, as this phenomenon is observed in 77% of the conserved exons.[50]

The VKORC1 gene is highly conserved across organisms and no south-east Asian house mouse from Browse Island carried mutations in the coding sequence that are linked to the *M. spretus* allele. The absence of these adaptive mutations is consistent with the absence of exposure to ARs.[13]

### Low genetic diversity of *Mus musculus* in Western Australia

*Mus musculus domesticus* of the Perth metropolitan area belongs to two closely related haplotypes that differ from each other by a single base pair change, and are most closely related to mice from European countries, such as The Netherlands and United Kingdom. A phylogeographic study of mice in Australia detected 12 haplotypes, with Western Australian house mice belonging to two major clades.[36] The clade occurring geographically closest to Perth is the most widespread clade in Australia, and this haplotype group is also the most common in the United Kingdom, which suggests Western Australian *M. m. domesticus* likely originated from the British Isles.[36]

Apart from mice from Madeira, which were grouped closely in the haplotype network, no countries were characterised by pockets of diversity. The resulting network was highly reticulate, indicating western house mice are closely related worldwide (Fig 1). The genotypic analysis of the mitochondrial D-loop of house mice from the Perth metropolitan area indicated the presence of a single, large panmictic population. Our network places Iranian mice in a central position (Fig 1), indicating Iran is an old (or ancestral) population[52] and potentially a centre of origin for the subspecies. This concurs with the assumed origin of the commensal relationship between mice and humans that was established in the Middle East around 10,000 BC when humans first formed large settlements in the Fertile Crescent.[53] The expansion of *M. m. domesticus* would have been aided by human transport and trade into the Mediterranean Sea during the Iron Age, and then to colonies like Australia by European settlers. [54–56]

A previous phylogeographic study of *M. musculus* in Australia documented the presence of *domesticus* as the predominant and only subspecies of house mouse in Australia[36], however mitochondrial D-loop data from mice of Browse Island from our study confirm previous records based on morphology suggesting that the *castaneus* subspecies of house mouse is present in Australia. However, unlike the European origins of the house mouse in Australia, the haplotype network of *M. m. castaneus* suggests that the origin of these Browse Island mice is China and Indonesia. Previous studies have also identified the Indonesian islands of Rote and Timor as the source of Australian *M. m. castaneus*.[57]

### Implications for pest management

When planning pest eradications or population control, it is vital to gather information on the resistance of the population in order to choose a control method that can successfully eliminate most of the population,^70^ ideally 99% of the population.[31] However, eradication attempts and rodent control programs in Western Australia have been carried out with no knowledge of the level of resistance in rodent populations.

As resistance-conferring VKORC1 mutations were not detected in the Perth metropolitan area, nor on Browse Island, house mouse populations in these areas can be managed with the use of a) FGARs with fewer negative impacts on non-target species or b) weaker SGARs when the natural tolerance exhibited by mice poses restrictions on the use of first generation compounds. Even so, regular testing should be used to detect developing resistance which can then be eliminated by a temporary switch to a SGAR,[33] or methods with low risk of bioaccumulation and secondary toxicity such as lethal trapping or a non-anticoagulant rodenticide containing actives such as cholecalciferol or corn gluten meal.[9,58]

As Perth is close to a major commercial port,[59] the risk of resistant mice invading the area through international shipping is high. Despite a reported genetic resilience of established mouse populations to new migrants,[60] in the event that resistant mice reach Perth these new individuals would be at a highly advantageous position due to the intense use of ARs.[14] Although our data indicate that, so far this has not occurred, it is an ongoing risk. A biosecurity and quarantine system has been developed to defend Western Australia and all other States and Territories on the Australia continent against incursions of pests, and checkpoints are in place at the State borders and all airports[61,62] and stringent biosecurity protocols need to be established for all ports also.[63] Vessels should be certified free of rodents and provide evidence that no re-infestation occurred since the certificate was issued.[63] Furthermore, resistance to ARs is not always concomitant in rat and mouse species, and if rats and mice of Perth exhibit different levels of resistance, more complex control programs are needed.[33]

The presence of a different mouse subspecies on Browse Island poses a biosecurity risk to mainland Australia.[64] Fortunately, the introgression between Browse Island mice and *M. spretus* was limited to intronic sequences in the VKORC1 gene, and resistance-related mutations were not carried by any individual from Browse Island that has been sequenced. Since the south-east Asian house mouse population on Browse Island carries silent VKORC1 mutations that do not impair rodenticide activity, the use of ARs to achieve pest eradication is possible, reducing the biosecurity risk of invasion of mainland Australia and will promote breeding of indigenous species by removing the risk of egg predation of both seabirds and green turtles (*Chelonia midas*) that nest on the island[1],[57]. The use of anticoagulant rodenticides on all islands should be carefully planned to reduce the impact of toxicants to endemic fauna of the island and coastal waters, since broadcast applications of rodenticides on islands are linked to exposure of non-target wildlife in coastal environments.[65] The waters around Browse Island are part of the Australian Fishing Zone, and are often visited by Indonesian fishermen who harvest marine organisms for consumption; as this biota might be exposed to ARs, the use of toxicants threatens human health through possible secondary exposure.[65,66]

## Conclusion

The order Rodentia is characterised by a high number of successful invaders, which humans have attempted to manage with the use of anticoagulant rodenticides (ARs) since the 1940s.[4,7] However, this has selected for resistance in the rodents and negatively impacted non-target species.[7,9,22,24] VKORC1 screening in multiple European countries has detected high frequencies of mutations that confer resistance to ARs.[11,18,19,39] In Australia, strong ARs can be purchased from supermarket and retail outlets and are routinely and intensively used in residential and commercial settings.[9] These factors led to the hypothesis that *Mus musculus* of Perth have a high probability of carrying resistance-related mutations in the VKORC1 gene. The aim of this study was to provide the first genetic data on AR resistance levels in two contrasting house mouse populations, and to make suggestions for conservation and biosecurity based on the evidence gathered.

Concomitantly this is the first genetic data available for resistance in mice on the Australian continent. The absence of resistance-conferring VKORC1 mutations in both Perth metropolitan area and Browse Island house mouse populations contrasts starkly with all other mouse populations screened thus far. Our results indicate that FGARs can be employed to control and/or eradicate mice effectively, minimising potential harm to non-target species, promoting biodiversity on the island[57] and reducing the biosecurity threat to mainland Australia posed by *Mus musculus castaneus*.

A general lack of data regarding the levels of AR resistance in commensal rodents in Australia and the impact of rodenticides on non-target biota should motivate future research to investigate resistance in mouse and rat populations throughout Australia. Investigation of genetic resistance should also include native rodents targeted for control in agricultural settings like *Rattus sordidus* and *Melomys burtoni*.[67] Anticoagulant rodenticides are likely to continue to be effective in controlling house mouse populations (at least in the short-term) and there should be a focus on: preventing the introduction of potentially resistant house mice to Western Australia through the port of Fremantle and other routes; genetic screening of other mainland Australian mouse and also rat populations; screening populations prior to eradication programs on islands to minimise the risk of coastal water contamination; and, generally reducing the threat of secondary poisoning to non-target species in both environments by careful planning and application of ARs underpinned by a thorough understanding of resistance in the target populations.

## Acknowledgements

We thank Dr J Lo and Ms K Patak for statistical advice. The authors thank the staff of the Department of Biodiversity, Conservation and Attractions for donating house mouse samples and for their useful insights into the history of human movements on Browse Island. We also thank the many volunteers across the Perth metropolitan area who donated house mouse samples to this project.

